# iModulonDB: a knowledgebase of microbial transcriptional regulation derived from machine learning

**DOI:** 10.1101/2020.08.13.250159

**Authors:** Kevin Rychel, Katherine Decker, Anand V Sastry, Patrick V Phaneuf, Saugat Poudel, Bernhard O Palsson

## Abstract

Independent component analysis (ICA) of bacterial transcriptomes has emerged as a powerful tool for obtaining co-regulated, independently-modulated gene sets (iModulons), inferring their activities across a range of conditions, and enabling their association to known genetic regulators. By grouping and analyzing genes based on observations from big data alone, iModulons can provide a novel perspective into how the composition of the transcriptome adapts to environmental conditions. Here, we present iModulonDB (imodulondb.org), a knowledgebase of prokaryotic transcriptional regulation computed from high-quality transcriptomic datasets using ICA. Users select an organism from the home page and then search or browse the curated iModulons that make up its transcriptome. Each iModulon and gene has its own interactive dashboard, featuring plots and tables with clickable, hoverable, and downloadable features. This site enhances research by presenting scientists of all backgrounds with co-expressed gene sets and their activity levels, which lead to improved understanding of regulator-gene relationships, discovery of transcription factors, and the elucidation of unexpected relationships between conditions and genetic regulatory activity. The current release of iModulonDB covers three organisms (*E. coli, S. aureus*, and *B. subtilis*) with 204 iModulons, and can be expanded to cover many additional organisms.

## INTRODUCTION

The transcriptional regulatory network (TRN) governs gene expression in response to environmental stimuli, which is of fundamental interest in biology. The TRN functions by employing condition-responsive regulators, such as transcription factors (TFs), to regulate the transcription of genes. Understanding these regulators and their effects in bacteria informs many important applications, including bioproduction (1) and antibiotic resistance (2). There are several organism-specific databases of gene-regulator relationships (3–5), but knowledge of regulator binding alone is insufficient to explain the complex responses reflected in transcriptomic datasets (6, 7). Thus, there is a need for data-driven approaches to TRN elucidation, which can detect the most important transcriptional signals in a gene expression dataset, identify their major gene constituents, and quantify their condition-dependent activity levels.

With the growing availability of bacterial transcriptomes, machine learning is emerging as a powerful tool for TRN elucidation. The falling price of RNA sequencing has led to a rapid growth in online transcriptomic databases (8, 9), creating a strong need for the development of analytical tools that can harness its scale to transform raw data into biologically meaningful information (10). For transcriptomic data, this knowledge comes in the form of 1) identifying which regulons are active in each condition probed in the dataset, 2) generating hypotheses about gene function and regulation, and 3) revealing novel relationships and patterns in bacterial lifestyles. In comparison, traditional methods such as chromatin immunoprecipitation (ChIP) assays (11), can be time-consuming and expensive, making them cumbersome for high-throughput discovery or hypothesis generation. They also do not yield the condition-specific strength of binding, which can be inferred by machine learning. Another strength of data-driven approaches is that they can be applied to any organism, regardless of prior information. Ultimately, a comprehensive, quantitative TRN would be the result of this pursuit.

Independent component analysis (ICA) addresses the goals described above. It is a blind source separation algorithm that identifies statistically independent signals underlying a dataset, and decomposes the original matrix into source (or module, **M**) and activity (**A**) matrices (12, 13). The module matrix **M** defines relationships between genes and the identified signals, while the activity matrix **A** describes the intensity of each signal in each sample. A comparison of 42 TRN inference methods demonstrated that ICA was the best at recovering known gene modules (14). Additional studies found that ICA-derived gene modules were robust across datasets (15). It has been used for a variety of organisms in the past, especially yeast and human cancer cells (16–20).

We recently applied ICA to high quality transcriptomic datasets for three species of bacteria: *Escherichia coli* (21), *Staphylococcus aureus* (22), and *Bacillus subtilis* (23). In our work with *E. coli*, we termed the independent signals “iModulons”, for **i**ndependently **modul**ated gene sets. We used the same codebase (www.github.com/SBRG/precise-db) to generate all three transcriptome decompositions. We then assigned categories, functions, and regulators to each iModulon. The regulator assignments were based on existing knowledge of transcription factor binding sites (3–5, 22), or, in some cases, a boolean combination of several regulons. By comparing our decompositions to known regulons described in other databases, we were able to identify potentially mislabeled or previously unknown TF-gene associations and verify them with ChIP-exo binding profiles. For example, we discovered five new binding sites for MetJ in *E. coli* (21). The observed iModulon activity levels also provided additional evidence for newly discovered regulatory roles or point to new hypotheses. For instance, we observed that the iron chelator pulcherrimin was active in stationary phase signaling in *B. subtilis* (23). This observation was recently externally validated (24). More broadly, each decomposition explained 65-80% of the variance in the transcriptomic data sets, indicating that the functions and regulators identified captured most of the transcriptional functions of the TRNs under the conditions where the data was obtained.

Several other studies have utilized iModulons to obtain valuable results. Our original *E. coli* study led to the discovery of a regulon putatively controlled by the uncharacterized TF *ydhC* (21), which has since been characterized and renamed (25). iModulons were also used to characterize the function of the TF OxyR (26), and to quantify the stress responses to heterologous gene expression (27). Additionally, iModulons captured the transcriptional effect of genomic alterations in adaptive laboratory evolution to naphthoquinone-based aerobic respiration (28).

In previous publications, curated iModulon dashboards were presented as static supplemental PDFs. While these are useful for disseminating basic information on specific iModulons, they suffered from several problems: 1) labeling of genes and inclusion of tables were limited by page space; 2) the inability to interact via hovering and clicking slowed analysis; 3) information was static and could not be updated; 4) access to the underlying data was limited and required coding experience; and 5) dissemination required sharing large PDFs instead of simple URL links. Given the potential for iModulons to enhance a wide range of research, we sought to address these issues with an online knowledgebase.

Here, we present the iModulon database (iModulonDB; imodulondb.org), an interactive web tool for accessing high quality transcriptomic datasets and their curated iModulons. Users begin by selecting their organism and dataset of interest, and then may either browse the iModulons identified through curated tables or search for the genes and regulators of interest to them. Each gene and iModulon has an interactive analytics dashboard featuring hoverable, clickable, and downloadable tables and graphs. iModulonDB presents the relationships between genes and iModulons, the inferred activities of regulators across diverse conditions, and the concordance between our data-driven gene modules and literature-defined regulons, such as those available on RegulonDB (3). Compared to the previous PDF dashboards, the new website enables global usage of this information and provides much more depth. iModulonDB meets the emerging need for an online, data-driven TRN resource; it obtains co-regulated gene sets based only on transcriptomic observations, which will help a broad audience of microbiologists and systems biologists to investigate genes or iModulons of interest.

## MATERIALS AND METHODS

### Data Generation and Acquisition

*E. coli* PRECISE (21) and *Staph*PRECISE (22) were generated using RNA sequencing (RNA-Seq), as described in their respective publications. The *Bacillus subtilis* dataset (23) is a well-known microarray dataset originally published by Nicolas, *et al*. (29), and is already featured as the expression compendium on the popular database *Subti*Wiki (4). Despite using older, established data, the *Bacillus* decomposition still provides novel insights, which demonstrates the efficacy of ICA. All raw data is available on GEO or SRA.

### Quality Control and Preprocessing

Prior to running ICA, we ensure that transcriptomic data passes stringent quality control as described in each dataset’s original publication (21–23). For RNA-Seq data, genes shorter than 100 nucleotides or with under 10 fragments per million-mapped reads are removed. We then compute transcripts per million (TPM) using DESeq2 (30). The final expression compendium is log-transformed log_2_(TPM + 1) before analysis – this is the form of the data available for download on iModulonDB. For both RNA-Seq and microarray data, biological replicates with R^2^ < 0.9 between final expression values are removed to reduce technical noise. Before computing iModulons, we choose a reference condition (such as exponential growth on M9 minimal media) and subtract its expression from all other samples; this results in activity levels for each iModulon relative to a known baseline.

### Computing Robust iModulons

We perform ICA as described in the original publications (21–23) using the Scikit-learn (31) implementation of the FastICA algorithm (32). To ensure robustness (since ICA is a stochastic gradient search algorithm), we perform ICA multiple times with random seeds for each dataset and cluster their **M** matrices using the Scikit-learn implementation of the DBSCAN algorithm (33). We keep independent components that appear in more than 50% of the ICA runs. Code to compute robust independent components is publicly available (github.com/SBRG/precise-db). Note that previous publications used **S** to refer to the **M** matrix, but it has since been renamed to avoid confusion with the stoichiometric matrix **S** (34).

ICA produces a set of independent components, each of which contains a weight for each gene in the expression dataset. Most gene weights are near zero for a given independent component, so the genes with large positive or negative weights are considered to be in the iModulon. To transform an independent component into an iModulon, we iteratively remove the highest absolute weighted genes from an iModulon until the D’Agostino K^2^ statistic of normality (35) of the remaining distribution falls below an organism-specific cutoff, and take all removed genes to be iModulon members. See the original publications for additional details (21–23).

## RESULTS

### A web-based analytics platform for data-driven TRNs

We developed iModulonDB (imodulondb.org) to enhance the field of microbial genetic regulation by presenting TRNs based on observed signals in transcriptomic datasets. This site provides biologists with the ability to easily navigate large datasets and quickly find gene modules through a search tool or by browsing curated annotations. The inferred activity levels of each iModulon, which are readily available on the site, provide valuable, novel insights into cellular function. We hope that iModulonDB will become an important part of the database ecosystem, providing a machine learning-derived perspective that also links to other databases for synergetic TRN characterization. **Figure 1** demonstrates the relationship between the content of an iModulon page (bottom left) and the original decomposition (top), as well as the types of insights that may be gleaned from this analysis (bottom right).

**Figure 1.**
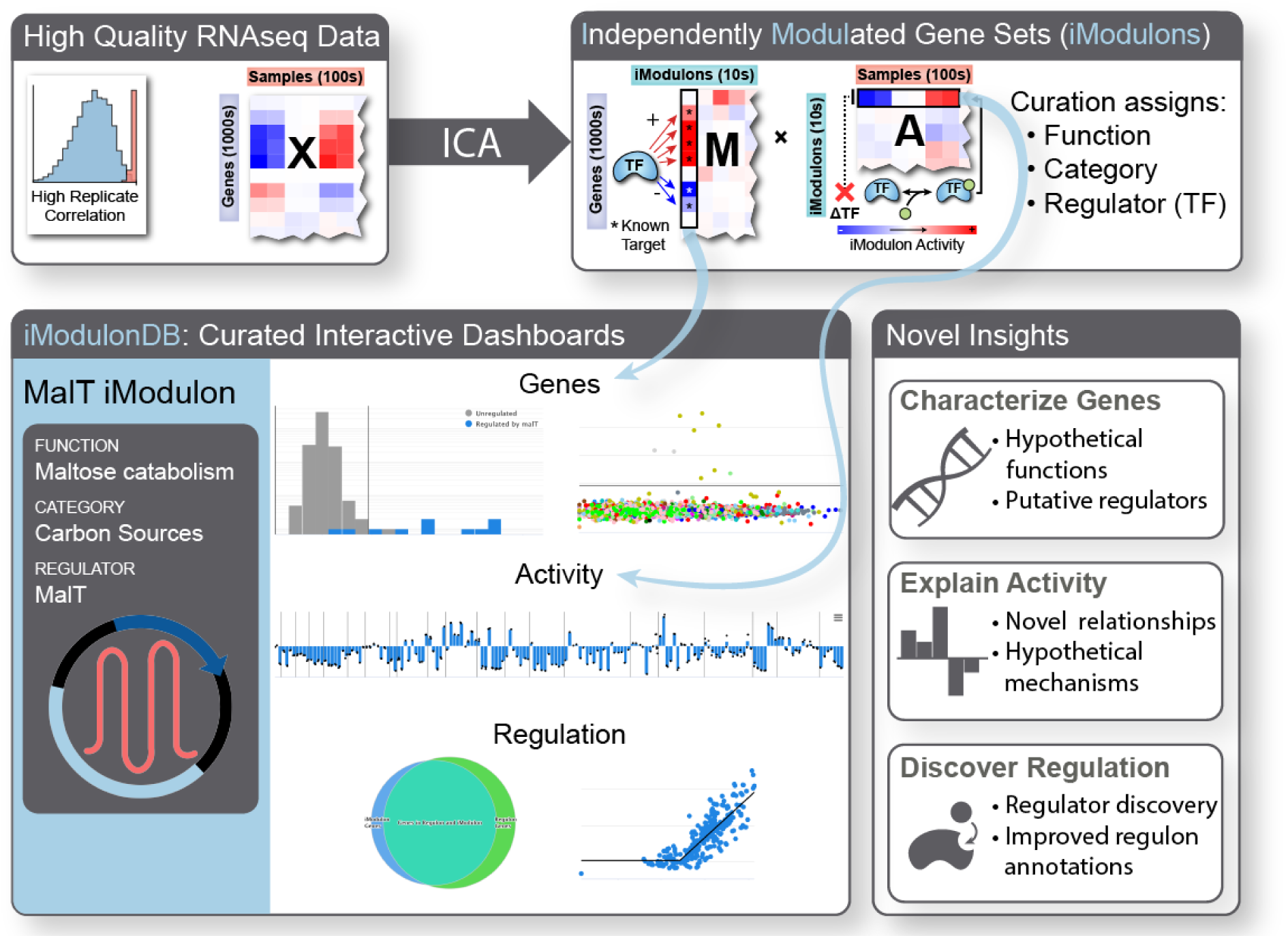
General outline of iModulonDB pipeline. Analysis begins with RNA-Seq data **X** (upper left), which is quality controlled for high biological replicate correlation. ICA is used to obtain iModulons, which have gene weights for each gene in the matrix **M** and activity levels for each condition in the matrix **A** (upper right). **M** weights are analogous to the strength of transcription factor binding upstream of genes and **A** activities are analogous to condition-dependent transcription factor activities, which may depend on processes such as ligand binding. Note that **X**≈**M*A**. The iModulons are curated by assigning functions, categories, and regulators, as shown in the example MalT iModulon dashboard. A representative screenshot of the interactive graphs is shown (middle; larger version with labels included in **Figure 2**), covering genes and activities as well as the concordance between this gene set and its curated regulator. The three rows of the dashboard result in the three categories of novel insights described (bottom).

iModulonDB currently contains the three datasets listed in **Table 1**. It covers 204 iModulons, 180 of which are characterized. The *E. coli* dataset is the largest; it includes 278 expression profiles with various gene knock-outs and evolved strains. This leads to its high dimensionality and lowest explained variance. The presence of genomic alterations also results in the most genomic iModulons (36), reducing the fraction of iModulons with known regulators. The *S. aureus* dataset is comparatively smaller, and is explained very well by its iModulons. Nearly all iModulons could be characterized, but the dearth of existing TRN annotations for this organism makes it more difficult to align certain iModulons to regulators. For *B. subtilis*, both mRNAs and non-coding RNAs were included in the decomposition. This leads to a high number of “genes” and 17 uncharacterized (noisy) iModulons. However, 95% of characterized iModulons were associated with a known set of regulators. Overall, the decompositions successfully produced iModulons that align well with our existing knowledge of transcriptional regulation in these species.

**Table 1.** Representative statistics of the three datasets currently in iModulonDB. ‘Genes’ and ‘Samples’ refer to the dataset size. Note that in *B. subtilis*, non-coding RNAs were included in the tiling array, which leads to the high gene count. ‘Conditions’ are unique experimental conditions, not counting biological replicate samples. ‘Dimensionality’ is the number of orthogonal principal components needed to explain 99% of the variance in the data (**Supplementary Methods**). The ‘Characterized iModulons’ row lists values and percentages of iModulons that can be categorized and named (excluded iModulons may result from noise or contain genes for which little information is available). ‘iModulons with Regulators’ are the subset of characterized iModulons that are mapped to regulators (excluded iModulons may be effects of genomic alterations or biological enrichments with no known regulators). ‘Explained Variance’ is the fraction of variance explained by reconstructing the original matrix using only iModulon member genes and activities (**Supplementary Methods**).

|  | *E. coli* (21) | *S. aureus* (22) | *B. subtilis* (23) | Total |
| --- | --- | --- | --- | --- |
| **Dataset Description** | RNA sequencing at SBRG | RNA sequencing at SBRG | Tiling Microarray, Nicolas, *et al.* (29) |  |
| **Genes** | 3923 | 2820 | 5875 | 12618 |
| **Samples** | 278 | 108 | 265 | 651 |
| **Conditions** | 163 | 54 | 104 | 321 |
| **Dimensionality** | 200 | 73 | 95 | 368 |
| **Total iModulons** | 92 | 29 | 83 | 204 |
| **Characterized iModulons** | 86 (93.5%) | 28 (96.6%) | 66 (79.5%) | 180 (88.2%) |
| **iModulons with Regulators** | 58 (63.0%) | 19 (65.5%) | 63 (75.9%) | 140 (68.6%) |
| **Explained Variance** | 68% | 76% | 72% |  |

### Dataset pages show lists of iModulons, constituting a data-driven TRN

Users begin by selecting one of our decompositions from the home page. This brings them to a dataset page (**Fig. 2A-B**). The left sidebar (**Fig. 2A**) lists basic features of the dataset, including organism details, dimensions, and a link to the relevant publication. The page also contains a curated table of all the iModulons identified (**Fig. 2B**). iModulons are given a name, regulator, function, and category. When multiple regulators capture the iModulon better than a single one, they may be combined using “or” (/) or “and” (+) set operators. For example, the ExuR/FucR iModulon contains genes regulated by either ExuR or FucR, while the GutM+SrlR iModulon contains only genes regulated by both GutM and SrlR (see the original publications (21–23) for more details). If multiple iModulons are linked to the same regulator, which indicates subsets of a large regulon, then a hyphen and numeral is added to the iModulon name (e.g. Fur-1). Some iModulons are enriched for a function or genomic alteration instead of a regulator, so they are named accordingly. Three statistics are also included in the table: 1) N, the number of genes; 2) precision, the fraction of iModulon genes regulated by the enriched regulator(s); and 3) recall, the fraction of regulated genes in the iModulon. Users may sort the table by any of its columns, or click on a row to learn about the iModulon.

**Figure 2.**
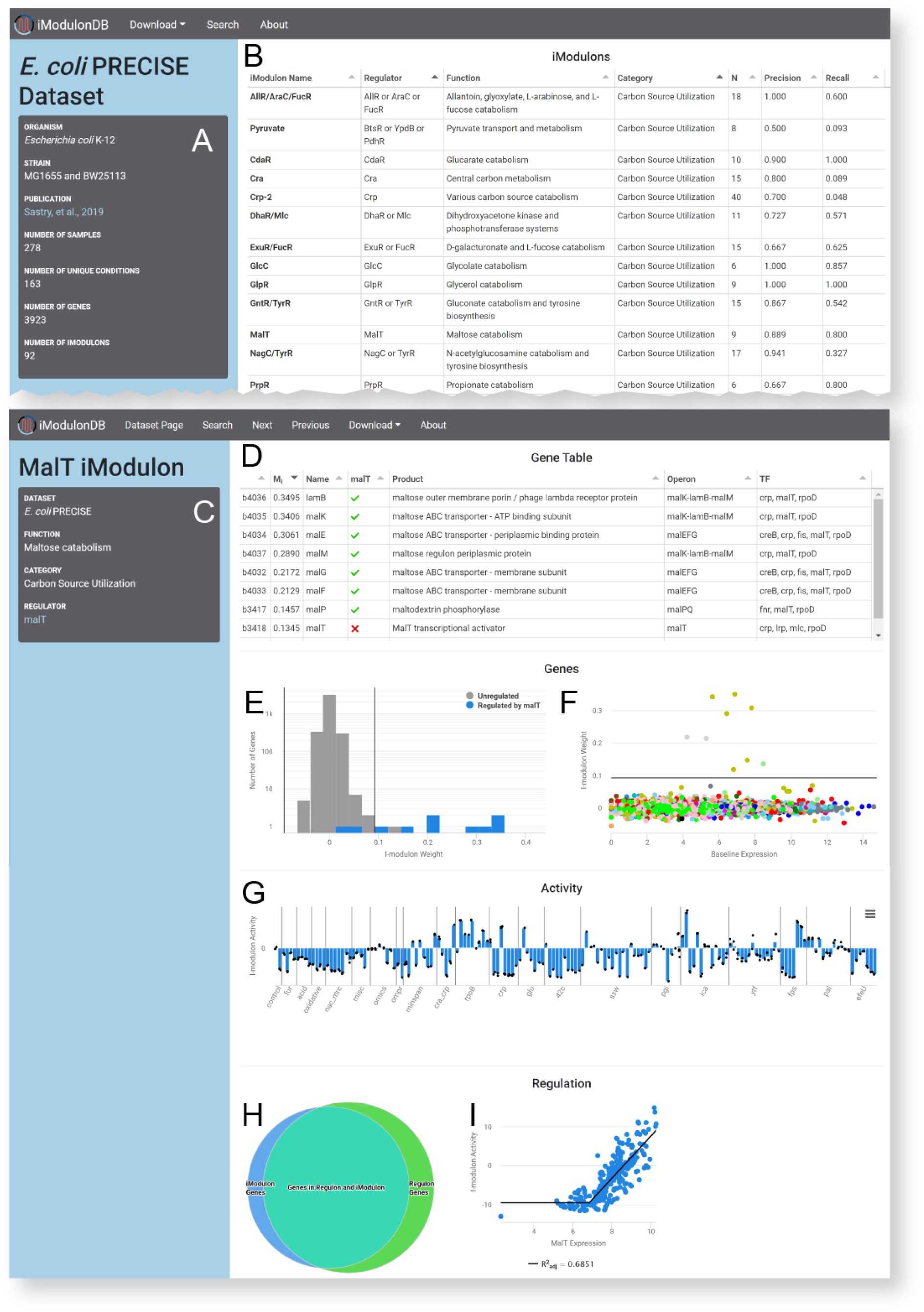
Representative screenshots of dataset and iModulon pages. **A-B**. Dataset page for *E. coli* (https://imodulondb.org/dataset.html?organism=e_coli&dataset=precise1). **A**. Details of the dataset, including a link to the relevant publication. **B**. List of iModulons for this dataset, along with some curated details and statistics. Click a row to access the appropriate iModulon page. Screenshot includes the first 13 iModulons on the table, which all happen to be in the category ‘Carbon Source Utilization’. **C-I**. iModulon page for the MalT iModulon in *E. coli* (https://imodulondb.org/iModulon.html?organism=e_coli&dataset=precise1&k=12). **C**. Details, including a link to its regulator’s page on RegulonDB. **D**. Gene table, listing the members of this iModulon and their annotations, plus whether or not they are annotated as being regulated by the iModulon’s regulator, MalT. **E**. Histogram showing the distribution of gene weights. Vertical bars represent the thresholds between member genes and non-member genes. All member genes are positively weighted in this example. **F**. Scatter plot of gene weights versus their expression in the baseline condition, color coded by COG. The horizontal line represents the weight threshold. **G**. Activity bar graph for this iModulon. Vertical lines separate projects within the dataset, bars represent mean activity of conditions, and individual black dots represent samples. **H-I**. This row only exists if the iModulon has a curated regulator. **H**. Venn diagram comparing this set of iModulon genes (left) to the set of genes annotated as being regulated by MalT (right). **I**. Scatter plot comparing the expression of MalT to the iModulon activity for all conditions. A broken line is used to indicate that there is a minimum expression of MalT before the correlation is observed.

Some common iModulon categories are: carbon source utilization, amino acid and nucleotide biosynthesis, metal homeostasis, stress response, and lifestyle (such as biofilm production). There are also iModulons categorized as ‘uncharacterized’, which may indicate undiscovered genetic or regulatory relationships (36).

### iModulon pages present information-dense interactive analytic dashboards

After selecting an iModulon, users are taken to its iModulon page (**Fig. 2C-I**). A gray box on this page (**Fig. 2C**) lists the curated features: function, category, and regulator. If available, the regulator entry will contain links to other databases (RegulonDB in *E. coli* (3) or *Subti*Wiki in *B. subtilis* (4)). If a regulator is assigned, the precision and recall of the enrichment is shown.

The gene table (**Fig. 2D**) contains information about all gene members of the iModulon. By default, the gene table is sorted by the associated gene weight from the **M** matrix (although users may click any column to sort by a different feature). Most columns contain information from external databases (EcoCyc in *E. coli* (37), AureoWiki in *S. aureus* (38), and *Subti*Wiki in *B. subtilis* (4)), such as gene product descriptions, operons, and associated regulators. If the iModulon has been assigned regulator(s), then the regulator names will appear as columns in the table, with green checks if the gene is known to be associated with that regulator and red X’s if not. This table is valuable for understanding the iModulon, as browsing the gene function list helps to understand the function of the iModulon as a whole. The boolean transcription factor columns are also a tool for discovery, since the genes marked with red ‘X’s may be controlled by the regulator despite the lack of a known association. Rows are also links to gene pages.

The gene histogram (**Fig. 2E**) shows the distribution of all gene weights for this iModulon. The gene scatter plot (**Fig. 2F**) presents the weights on the Y axis and the expression of each gene in the reference condition along the X axis. For details of the features and analysis of these two plots, see the supplement (**Supplementary Results**).

The activity bar graph (**Fig. 2G**) displays the activity level of the iModulon (with respect to a reference condition) across all expression profiles. iModulon activities are a measure of relative transcription factor activity, which can be a very valuable resource that is difficult to obtain using other methods.

If the iModulon has a matched regulator, then a ‘Regulation’ row will appear at the bottom of the dashboard (**Fig. 2H-I**), which quantifies the concordance between our transcriptomics-derived groupings (iModulons) and the gene groupings (regulons) available in literature and on other databases (3, 4). It contains a venn diagram comparing the set of iModulon genes against the regulon (**Fig. 2H**), which can be hovered over to quickly see agreements and discrepancies between iModulons and regulon annotations. If the regulator is also encoded by a gene in the expression compendium, then there will be a scatter plot or series of scatter plots showing the iModulon activity and regulator gene expression across the conditions (**Fig. 2I**). Each point can be hovered over to see the name of the sample it represents. For some regulators, such as sigma factors and self-regulating TFs, a high correlation may be observed in this plot. When post-transcriptional modifications such as phosphorylation and ligand binding affect the iModulon activity, we see low correlations between the expression of the regulator and the activity of the genes it regulates. Regulators are not necessarily part of their own iModulons. The computation of these correlations is described in the original dataset publications (21–23). In some cases, the correlation is best captured using a broken line (as in **Fig. 2I**) to signify that a minimum expression level of the regulator must be reached before a correlation is observed.

### Gene pages connect users to iModulons of interest

Gene pages enable users to quickly find iModulons relevant to their research. Similar to the iModulon and dataset pages, the left sidebar on this page will list the gene’s identifier, gene name, gene product, operon, COG category, and known regulator(s), as well as a link to a relevant database (EcoCyc in *E. coli* (37), Aureowiki in *S. aureus* (38), and *Subti*Wiki in *B. subtilis* (4)). The dashboard contains two elements: 1) a table of iModulons and 2) an expression profile. The expression profile shows log_2_ TPM expression values across each dataset.

The iModulon table (included in **Fig. 3B**) lists the iModulons in order of weight, with the strongest iModulon associations at the top of the list. If the gene is in an iModulon, a green check will appear next to it (red X otherwise). The table also includes additional details for each iModulon. Here, a user can easily find iModulons containing genes of interest, and navigate to the iModulon page to learn more. If a gene is not in any iModulon, the gene is not a strong part of any independent signals in the dataset.

**Figure 3.**
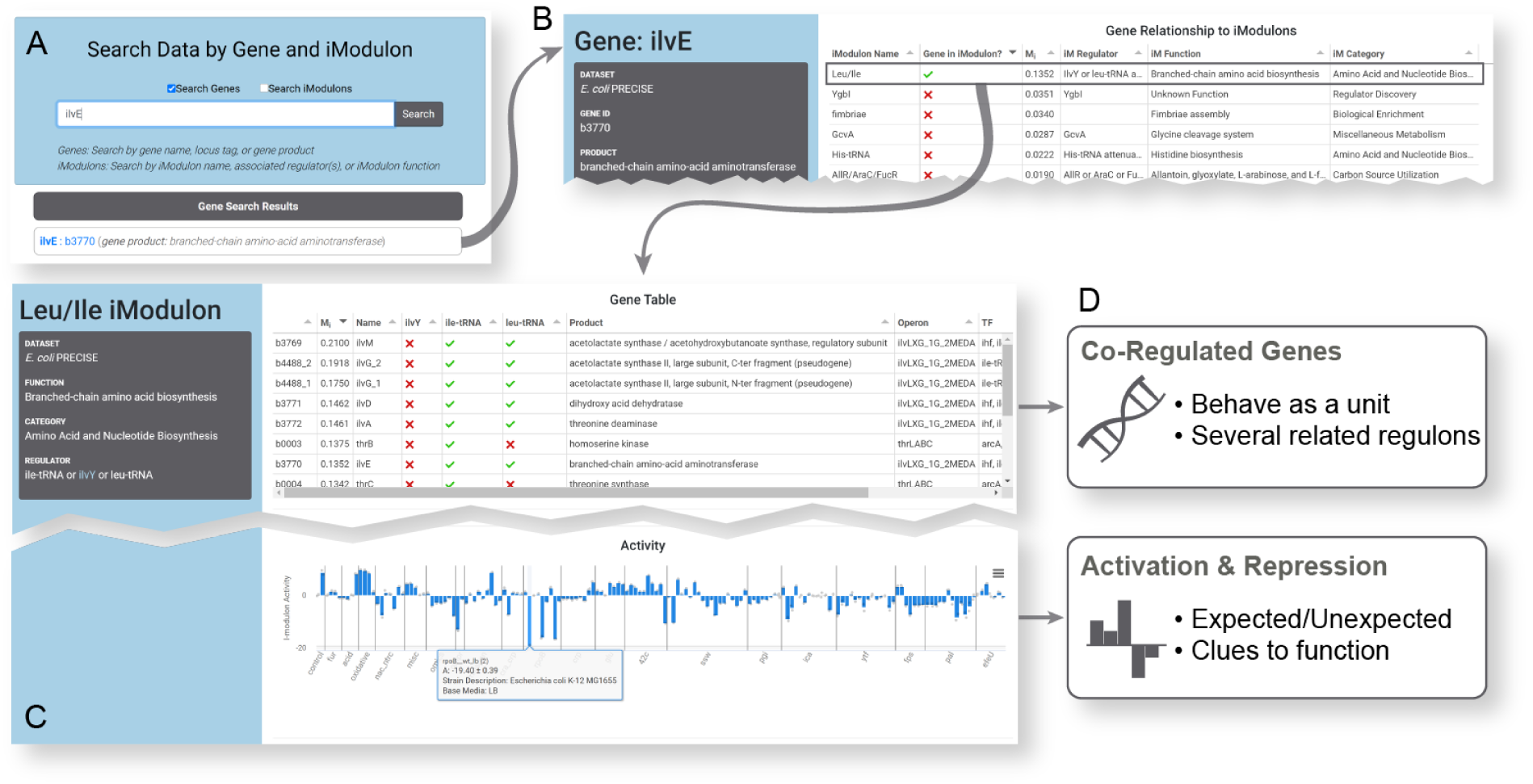
Flowchart of an example analysis for the gene *ilvE*. **A**. The user searches for their gene of interest, *ilvE*. This returns the gene as a result, which can be clicked to access its gene page (**B**). **B**. The iModulon table on the gene page shows that this gene is the member of the Leu/Ile iModulon. Clicking that row in the table brings the user to the iModulon page (**C**). **C**. Two features of the iModulon page are shown: the gene table and activity bar graph (see **Figure 2** for details). The activity bar graph includes an example tooltip, which appears when the bar is hovered over with a mouse. In this case, the bar with the tooltip is a repressing condition labeled with LB media, which is expected to repress the function of this iModulon. **D**. The insights gained are summarized in the boxes: the co-regulated genes are enumerated, and the activating or repressing conditions in our dataset can be determined.

### Other major features: About, Search and Download

As many researchers are not familiar with iModulons, we have included a thorough “About” page with a YouTube video on ICA, an illustrated walkthrough of our pipeline, and details on how to read our dashboards. We also include an email address where users can send feedback or request that we work with their dataset.

After choosing a dataset, users may click the “Search” link in the toolbar from any page to access our search functionality. This tool allows the users to search through genes and/or iModulons. For genes, results matching queried gene names, gene IDs, and gene products will be displayed. Similarly for iModulons, matching iModulon names, associated regulators, and iModulon functions will generate search results. Results are separated into the relevant iModulon and gene sections.

Each of the main three pages (dataset, iModulon, and gene) has a download menu in the toolbar. From there, all data is available for download in bulk or parts. The bulk data includes the original log-tpm data, the two ICA-generated matrices, the gene annotations and literature TRN used, the metadata on all samples, and the curated iModulon table. For individual iModulons, the gene weights and activity levels are available along with the gene table as it appears in the dashboard. Likewise, gene pages support the download of all iModulon associations to that gene, its expression levels for each sample, and the iModulon table as it appears in the dashboard. We encourage data download and custom analysis.

### Case Study: How To Use iModulonDB to Enhance Research

This section includes an example of a researcher who could gain valuable information from iModulonDB, and how they might do so. **Figure 3** illustrates the process.

Researcher X has chosen a gene of interest. They are studying the branched chain amino acid (BCAA) synthesis enzyme encoded by *ilvE* in *E. coli* as a potential drug target. They want to know which conditions activate *ilvE* expression, and what other genes are co-expressed with this gene. Researcher X can navigate to the *E. coli* dataset on iModulonDB, and then choose “Search” in the toolbar and enter “ilvE” (**Fig. 3A**). *ilvE* will appear as a gene page result, which links to the *ilvE* gene page (**Fig. 3B**). From there, they can see some basic information about the gene, a link to its EcoCyc page (37), and its expression across our compendium. They will also find the iModulon table, which shows that it is a member of the “Leu/Ile” iModulon. As expected, the function of the iModulon is BCAA biosynthesis. Clicking the “Leu/Ile” row in the iModulon table will take them to the corresponding iModulon page (**Fig. 3C**) where they can see all of the genes (including *ilvE*) in this independent signal of the dataset, as well as its activity across all conditions. The iModulon is annotated as being regulated by ile-tRNA, ilvY, or leu-tRNA. IlvY is a transcription factor with a page on RegulonDB (3), so users can follow that link to learn more about its regulon.

Along with linking to relevant pages on other databases, iModulonDB provides Researcher X with some novel information (**Fig. 3D**). The first would be the particular gene grouping of this iModulon – it covers the synthesis of all three BCAAs plus threonine as a single unit of the transcriptome, instead of three separate regulons or several operons as they may find on other databases. This grouping is observed from transcriptomic data, and likely results from a combination of genetic factors that respond to separate but metabolically related metabolites. It is useful to understand the cell as manipulating all of these genes together as a unit, which may affect how the researcher implements their drug targeting. The other major insight would be gained by probing the activity profile; for example, hovering over the tallest bars shows that this iModulon is activated under iron starvation and reactive oxygen species (ROS) stress and hovering over the most negative bars indicates deactivation under rich media conditions or under osmotic stress from NaCl. Searching the literature for explanations would reveal that ROS and iron starvation damage the iron-sulfur clusters needed for member enzymes (39) and create BCAA-limiting conditions, while the amino acids in rich media turn off these genes through their well-studied mechanisms (40). Direct or indirect repression of these genes by osmotic stress has not been studied and may be worth investigating. All of this information would help Researcher X determine whether *ilvE* and the function encoded by the Leu/Ile iModulon are good potential drug targets.

Not all iModulons need to be found through gene pages. Users may be more interested in a general process or specific regulator, which can also be searched for on the search page. Alternatively, users can start by selecting an iModulon from the dataset page. They may stumble upon a surprising gene grouping or unexpected condition/iModulon activation pair, which could lead to valuable new hypotheses worth testing. This knowledgebase would also be very valuable for someone studying a completely uncharacterized gene, as its presence in an iModulon gives clues to its function. Another way to use this knowledgebase is simply to follow along with the existing publications (21–23). For *E. coli* and *B. subtilis*, expected and unexpected activating conditions for each iModulon were listed in supplementary tables, which may serve as a starting place for hypothesis generation.

### Design and Implementation

iModulonDB is implemented and deployed using a simple web application stack and combination of user interface technologies. The server side is entirely hosted by GitHub Pages (pages.github.com) with HTTPS enforced. It relies on local computations performed in Python 3.7 with Jupyter notebooks (jupyter.org). The client side is implemented using a combination of HTML, CSS, and JavaScript.

Bootstrap 4.5 (getbootstrap.com) is used to manage page layout and ensure mobile compatibility. Interactive tables were made using Tabulator (tabulator.info), and plots are implemented in HighCharts using a non-commercial license (highcharts.com). Other JavaScript packages used include: jQuery (jquery.com), Popper.js (popper.js.org), URLSearchParams (developer.mozilla.org), and PapaParse (papaparse.com).

## DISCUSSION

iModulonDB is a data-driven bacterial TRN knowledgebase that meets the need for unsupervised, observation-based gene groupings and inferred genetic regulator activities. Its interactive graphical user interface with search and download functionality facilitates usage by scientists with computational and non-computational backgrounds alike. The platform provides a novel perspective on bacterial genomes based on big data observations, and also connects to (and largely agrees with) databases that report established gene groupings based on past literature. It currently includes data for three organisms based on three publications and contains 204 iModulons, and will be expanded in the future to accommodate new datasets and organisms. This work demonstrates the potential for iModulonDB to improve our understanding of bacterial TRNs for a wide variety of organisms.

## AVAILABILITY

iModulonDB is freely available online at https://imodulondb.org and can be accessed with a JavaScript-enabled browser. The download links in the toolbars enable download of all data and facilitate custom analysis.

## SUPPLEMENTARY DATA

Supplementary Data are available at NAR online.

## ACKNOWLEDGEMENT

The authors would like to thank Dr. Akanksha Rajput and David Bour for their input on website infrastructure and code. We would also like to thank Dr. Hyungyu Lim, Dr. Sonal Choudhary, and Reo Yoo for testing the site and providing valuable suggestions.

## AUTHOR CONTRIBUTIONS

K.R. led site development and drafted the paper. K.R. and K.D. developed the website. A.V.S., P.V.P., and B.O.P. provided mentorship and guidance in design and implementation. A.V.S., S.P., and K.R. provided, analyzed, and reformatted data for the site. All participated in writing the paper.

## REFERENCES

1. Gu, Y., Xu, X., Wu, Y., Niu, T., Liu, Y., Li, J., Du, G. and Liu, L. (2018) Advances and prospects of Bacillus subtilis cellular factories: From rational design to industrial applications. Metabolic Engineering, 50, 109–121.

2. Gunn, J.S. (2008) The Salmonella PmrAB regulon: lipopolysaccharide modifications, antimicrobial peptide resistance and more. Trends in Microbiology, 16, 284–290.

3. Santos-Zavaleta, A., Salgado, H., Gama-Castro, S., Sánchez-Pérez, M., Gómez-Romero, L., Ledezma-Tejeida, D., García-Sotelo, J.S., Alquicira-Hernández, K., Muñiz-Rascado, L.J., Peña-Loredo, P., et al. (2019) RegulonDB v 10.5: tackling challenges to unify classic and high throughput knowledge of gene regulation in E. coli K-12. Nucleic Acids Res., 47, D212–D220.

4. Zhu, B. and Stülke, J. (2018) SubtiWiki in 2018: from genes and proteins to functional network annotation of the model organism Bacillus subtilis. Nucleic Acids Res., 46, D743–D748.

5. Novichkov, P.S., Kazakov, A.E., Ravcheev, D.A., Leyn, S.A., Kovaleva, G.Y., Sutormin, R.A., Kazanov, M.D., Riehl, W., Arkin, A.P., Dubchak, I., et al. (2013) RegPrecise 3.0--a resource for genome-scale exploration of transcriptional regulation in bacteria. BMC Genomics, 14, 745.

6. Larsen, S.J., Röttger, R., Schmidt, H.H.H.W. and Baumbach, J. (2019) E. coli gene regulatory networks are inconsistent with gene expression data. Nucleic Acids Res, 47, 85–92.

7. Fang, X., Sastry, A., Mih, N., Kim, D., Tan, J., Yurkovich, J.T., Lloyd, C.J., Gao, Y., Yang, L. and Palsson, B.O. (2017) Global transcriptional regulatory network for Escherichia coli robustly connects gene expression to transcription factor activities. PNAS, 10.1073/pnas.1702581114.

8. Barrett, T., Wilhite, S.E., Ledoux, P., Evangelista, C., Kim, I.F., Tomashevsky, M., Marshall, K.A., Phillippy, K.H., Sherman, P.M., Holko, M., et al. (2013) NCBI GEO: archive for functional genomics data sets—update. Nucleic Acids Res, 41, D991–D995.

9. Leinonen, R., Sugawara, H. and Shumway, M. (2011) The Sequence Read Archive. Nucleic Acids Res, 39, D19–D21.

10. Margolis, R., Derr, L., Dunn, M., Huerta, M., Larkin, J., Sheehan, J., Guyer, M. and Green, E.D. (2014) The National Institutes of Health’s Big Data to Knowledge (BD2K) initiative: capitalizing on biomedical big data. J Am Med Inform Assoc, 21, 957–958.

11. Rhee, H.S. and Pugh, B.F. (2012) ChIP-exo method for identifying genomic location of DNA-binding proteins with near-single-nucleotide accuracy. Curr Protoc Mol Biol, Chapter 21, Unit 21.24.

12. Comon, P. (1994) Independent component analysis, A new concept? Signal Processing, 36, 287–314.

13. Shlens, J. (2014) A Tutorial on Independent Component Analysis. 1404.2986 [cs, stat].

14. Saelens, W., Cannoodt, R. and Saeys, Y. (2018) A comprehensive evaluation of module detection methods for gene expression data. Nature Communications, 9.

15. Cantini, L., Kairov, U., Reyniès, A. de, Barillot, E., Radvanyi, F. and Zinovyev, A. (2019) Assessing reproducibility of matrix factorization methods in independent transcriptomes. Bioinformatics, 35, 4307.

16. Zhang, X.W., Yap, Y.L., Wei, D., Chen, F. and Danchin, A. (2005) Molecular diagnosis of human cancer type by gene expression profiles and independent component analysis. European Journal of Human Genetics, 13, 1303–1311.

17. Kong, W., Vanderburg, C.R., Gunshin, H., Rogers, J.T. and Huang, X. (2008) A review of independent component analysis application to microarray gene expression data. BioTechniques, 45, 501.

18. Engreitz, J.M., Daigle, B.J., Jr, Marshall, J.J. and Altman, R.B. (2010) Independent component analysis: mining microarray data for fundamental human gene expression modules. Journal of biomedical informatics, 43, 932.

19. Karczewski, K.J., Snyder, M., Altman, R.B. and Tatonetti, N.P. (2014) Coherent Functional Modules Improve Transcription Factor Target Identification, Cooperativity Prediction, and Disease Association. PLOS Genetics, 10, e1004122.

20. Sompairac, N., Nazarov, P.V., Czerwinska, U., Cantini, L., Biton, A., Molkenov, A., Zhumadilov, Z., Barillot, E., Radvanyi, F., Gorban, A., et al. (2019) Independent Component Analysis for Unraveling the Complexity of Cancer Omics Datasets. International Journal of Molecular Sciences, 20, 4414.

21. Sastry, A.V., Gao, Y., Szubin, R., Hefner, Y., Xu, S., Kim, D., Choudhary, K.S., Yang, L., King, Z.A. and Palsson, B.O. (2019) The Escherichia coli transcriptome mostly consists of independently regulated modules. Nat Commun, 10.

22. Poudel, S., Tsunemoto, H., Seif, Y., Sastry, A.V., Szubin, R., Xu, S., Machado, H., Olson, C.A., Anand, A., Pogliano, J., et al. (2020) Revealing 29 sets of independently modulated genes in Staphylococcus aureus, their regulators, and role in key physiological response. PNAS, 117, 17228–17239.

23. Rychel, K., Sastry, A.V. and Palsson, B.O. (2020) Machine learning uncovers independently regulated modules in the Bacillus subtilis transcriptome. bioRxiv, 10.1101/2020.04.26.062638.

24. Arnaouteli, S., Matoz-Fernandez, D.A., Porter, M., Kalamara, M., Abbott, J., MacPhee, C.E., Davidson, F.A. and Stanley-Wall, N.R. (2019) Pulcherrimin formation controls growth arrest of the Bacillus subtilis biofilm. PNAS, 116, 13553–13562.

25. Rodionova, I.A., Gao, Y., Sastry, A., Yoo, R., Rodionov, D.A., Saier, M.H. and Palsson, B.Ø. (2020) Synthesis of the novel transporter YdhC, is regulated by the YdhB transcription factor controlling adenosine and adenine uptake. bioRxiv, 10.1101/2020.05.03.074617.

26. Anand, A., Chen, K., Catoiu, E., Sastry, A.V., Olson, C.A., Sandberg, T.E., Seif, Y., Xu, S., Szubin, R., Yang, L., et al. (2020) OxyR Is a Convergent Target for Mutations Acquired during Adaptation to Oxidative Stress-Prone Metabolic States. Mol Biol Evol, 37, 660–667.

27. Tan, J., Sastry, A.V., Fremming, K.S., Bjørn, S.P., Hoffmeyer, A., Seo, S., Voldborg, B.G. and Palsson, B.O. (2020) Independent component analysis of E. coli’s transcriptome reveals the cellular processes that respond to heterologous gene expression. Metabolic Engineering, 10.1016/j.ymben.2020.07.002.

28. Anand, A., Chen, K., Yang, L., Sastry, A.V., Olson, C.A., Poudel, S., Seif, Y., Hefner, Y., Phaneuf, P.V., Xu, S., et al. (2019) Adaptive evolution reveals a tradeoff between growth rate and oxidative stress during naphthoquinone-based aerobic respiration. PNAS, 116, 25287–25292.

29. Nicolas, P., Mader, U., Dervyn, E., Rochat, T., Leduc, A., Pigeonneau, N., Bidnenko, E., Marchadier, E., Hoebeke, M., Aymerich, S., et al. (2012) Condition-Dependent Transcriptome Reveals High-Level Regulatory Architecture in Bacillus subtilis. Science, 335, 1103–1106.

30. Love, M.I., Huber, W. and Anders, S. (2014) Moderated estimation of fold change and dispersion for RNA-seq data with DESeq2. Genome Biol., 15, 550.

31. Pedregosa, F. Scikit-learn: Machine Learning in Python. MACHINE LEARNING IN PYTHON.

32. Hyvärinen, A. (1999) Fast and robust fixed-point algorithms for independent component analysis. IEEE Trans Neural Netw, 10, 626–634.

33. Ester, M., Kriegel, H.-P., Sander, J. and Xu, X. (1996) A density-based algorithm for discovering clusters in large spatial databases with noise. In Proceedings of the Second International Conference on Knowledge Discovery and Data Mining, KDD’96. AAAI Press, Portland, Oregon, pp. 226–231.

34. Orth, J.D., Thiele, I. and Palsson, B.Ø. (2010) What is flux balance analysis? Nat Biotechnol, 28, 245–248.

35. D’Agostino, R.B. and Belanger, A. (1990) A Suggestion for Using Powerful and Informative Tests of Normality. The American Statistician, 44, 316–321.

36. Sastry, A.V., Hu, A., Heckmann, D., Poudel, S., Kavvas, E. and Palsson, B.O. (2020) Matrix factorization recovers consistent regulatory signals from disparate datasets. bioRxiv, 10.1101/2020.04.26.061978.

37. Keseler, I.M., Mackie, A., Santos-Zavaleta, A., Billington, R., Bonavides-Martínez, C., Caspi, R., Fulcher, C., Gama-Castro, S., Kothari, A., Krummenacker, M., et al. (2017) The EcoCyc database: reflecting new knowledge about Escherichia coli K-12. Nucleic Acids Res, 45, D543–D550.

38. Fuchs, S., Mehlan, H., Bernhardt, J., Hennig, A., Michalik, S., Surmann, K., Pané-Farré, J., Giese, A., Weiss, S., Backert, L., et al. (2018) AureoWiki ? The repository of the Staphylococcus aureus research and annotation community. Int. J. Med. Microbiol., 308, 558–568.

39. Yang, L., Mih, N., Anand, A., Park, J.H., Tan, J., Yurkovich, J.T., Monk, J.M., Lloyd, C.J., Sandberg, T.E., Seo, S.W., et al. (2019) Cellular responses to reactive oxygen species are predicted from molecular mechanisms. Proc Natl Acad Sci U S A, 116, 14368–14373.

40. Brosnan, J.T. and Brosnan, M.E. (2006) Branched-chain amino acids: enzyme and substrate regulation. J. Nutr., 136, 207S–11S.

