## Supplemental Information for "iModulonDB: a knowledgebase of microbial transcriptional regulation derived from machine learning"

### iModulonDB Supplementary Information

#### SUPPLEMENTARY METHODS

##### Computation of Dimensionality and Explained Variance

To estimate the dimensionality of each expression dataset, we compute the number of principal components required to explain 99% of the variance. This is computed using the sci-kit learn implementation (1) of the principal component analysis algorithm.

Explained variance of the iModulon decompositions was computed as described in the original publications (2–4). Briefly, the **M** matrix was modified so that all non-member iModulon-gene associations were exactly zero. Then this **M** matrix is multiplied by the **A** matrix to reconstruct a prediction of the original values using only iModulons and their activities. The percent of the variance in the original matrix that this prediction accurately explains is reported.

#### SUPPLEMENTARY RESULTS

##### Plots on the iModulon page provide detailed analytics of gene weights

The gene histogram (**Fig. 2E**) shows the distribution of weights associating this iModulon with all genes.It has a typical shape with a high number of non-member genes near zero and a small number of member genes outside of the vertical threshold lines. All bars can be hovered over to see a list or number of genes. If the iModulon is assigned multiple regulators, the histogram is color-coded by regulon – regulated genes with weights near zero may have other regulators such that those genes are not part of this independent signal in the transcriptome. Each element in the legend can be clicked to show or hide regulated genes.

The gene scatter plot (**Fig. 2F**) places the weights on the Y axis and the expression of each gene from the reference condition along the X axis. If the expression is low, these genes are not important in the baseline condition (for example, alternative carbon sources have low expression when in the presence of glucose). Each point is color-coded by cluster of orthologous groups (COG; computed via eggNOG (5, 6)), hovering over a point displays additional information, and clicking takes you to the corresponding gene page. This plot allows you to visualize COG categories and view gene annotations for both member and non-member genes.

##### Features of the activity bar graph enable understanding of condition-specific behavior

The activity bar graph on the iModulon page presents iModulon activity under each condition (**Fig. 2G**). Bars represent conditions and dots represent individual samples within that condition. Hovering over a bar displays some relevant metadata about culture conditions, and clicking it may take you to a paper with more details if it exists (for example, the *E. coli* dataset contains expression profiles from multiple projects, and each project has its own paper). The graph can also be zoomed in and scrolled horizontally, or viewed in full screen mode.

#### SUPPLEMENTARY REFERENCES


1. Pedregosa,F. Scikit-learn: Machine Learning in Python. *MACHINE LEARNING IN PYTHON*.
2. Sastry,A.V., Gao,Y., Szubin,R., Hefner,Y., Xu,S., Kim,D., Choudhary,K.S., Yang,L., King,Z.A. and Palsson,B.O. (2019) The Escherichia coli transcriptome mostly consists of independently regulated modules. *Nat Commun*, **10**.
3. Poudel,S., Tsunemoto,H., Seif,Y., Sastry,A.V., Szubin,R., Xu,S., Machado,H., Olson,C.A., Anand,A., Pogliano,J., *et al.* (2020) Revealing 29 sets of independently modulated genes in Staphylococcus aureus, their regulators, and role in key physiological response. *PNAS*, **117**, 17228–17239.
4. Rychel,K., Sastry,A.V. and Palsson,B.O. (2020) Machine learning uncovers independently regulated modules in the Bacillus subtilis transcriptome. *bioRxiv*, 10.1101/2020.04.26.062638.
5. Huerta-Cepas,J., Forslund,K., Coelho,L.P., Szklarczyk,D., Jensen,L.J., von Mering,C. and Bork,P. (2017) Fast Genome-Wide Functional Annotation through Orthology Assignment by eggNOG-Mapper. *Mol Biol Evol*, **34**, 2115–2122.
6. Huerta-Cepas,J., Szklarczyk,D., Heller,D., Hernández-Plaza,A., Forslund,S.K., Cook,H., Mende,D.R., Letunic,I., Rattei,T., Jensen,L.J., *et al.* (2019) eggNOG 5.0: a hierarchical, functionally and phylogenetically annotated orthology resource based on 5090 organisms and 2502 viruses. *Nucleic Acids Res*, **47**, D309–D314.
